# Glyphosate as a driver of antimicrobial resistance evolution in bacteria

**DOI:** 10.64898/2026.05.11.724250

**Authors:** Tuomas Tall, Marjo Helander, Jaime Iranzo, Lyydia Leino, Miia J. Rainio, Eero Vesterinen, Kari Saikkonen, Irma Saloniemi, Suni A. Mathew, Pere Puigbò

## Abstract

Glyphosate, the world’s most widely used herbicide, targets the enzyme 5-enolpyruvylshikimate-3-phosphate synthase (EPSPS), which is conserved across plants and many bacteria. While its environmental effects are increasingly recognized, its role on antimicrobial resistance (AMR) remains incompletely understood. In particular, the link between intrinsic glyphosate sensitivity and AMR gene content or evolutionary dynamics has not been systematically explored. We examined the relationship between bacterial sensitivity to glyphosate, AMR profiles, and the evolution of AMR genes. We analyzed genome datasets from the human gut microbiota and the Alignable Tight Genomic Clusters (ATGC). EPSPS sequences were identified *via* BLAST and annotations and classified based on the intrinsic sensitivity to glyphosate using the EPSPSClass webserver. AMR genes, including associated drug classes and resistance mechanisms, were annotated using the Comprehensive Antibiotic Resistance Database (CARD). Across datasets, glyphosate-sensitive bacteria carried a greater diversity of AMR genes and mechanisms. In contrast, probabilistic modeling revealed that glyphosate-resistant bacteria accumulate AMR genes at significantly higher rates. Phylogenetic birth-and-death analyses and stochastic mapping further revealed elevated AMR gene gain, loss, expansion, and reduction in resistant strains. These results indicate a decoupling between AMR gene diversity and evolutionary dynamics: sensitive bacteria maintain more resistance genes, whereas resistant bacteria display accelerated AMR gene turnover. This suggests that glyphosate resistance is linked to increased genome dynamics, potentially enhancing bacteria’s adaptability under combined herbicide and antimicrobial pressures. Given glyphosate’s extensive agricultural use and potential human exposure, these findings highlight an underappreciated link between herbicide resistance and the evolution of AMR in bacterial populations.

## MAIN

Humans have exploited antimicrobial compounds for millennia, as evidenced by the 1550 BC Ebers papyrus describing moldy bread for treating infections [1]. The modern antibiotic era began with Paul Ehrlich’s Salvarsan in 1910, followed by sulfonamides in the 1930s and Alexander Fleming’s discovery of penicillin in 1929 [1]. Resistance emerged rapidly [2, 3], driving the development of semisynthetic β-lactams following Dorothy Hodgkin’s structural elucidation in 1945 [4]. Subsequent exploitation of soil actinomycetes by Selman Waksman ushered in the antibiotic “golden age” (1940–1960), during which most major antibiotic classes were discovered [5, 6].

Since then, the extensive use of antibiotics in medicine, agriculture, and animal production has accelerated the global rise of antimicrobial resistance (AMR) [7]. This is fuelled by rapid bacterial evolution, biofilms, and horizontal gene transfer, which together facilitate the emergence and dissemination of resistance determinants across microbial communities [8, 9]. The human gut microbiome itself constitutes a major reservoir of AMR genes [10]. However, the discovery of new antibiotic classes has slowed markedly since the 1970s [11, 12], and high-throughput screening efforts have yielded limited breakthroughs [13]. These constraints have prompted increasing interest in alternative antimicrobial strategies, including bacteriophages, antimicrobial peptides, CRISPR-based gene-editing tools, and predatory bacteria [14]. Collectively, this context underscores the need to understand how diverse environmental pressures, beyond antibiotics, shape the evolution and spread of AMR.

A growing body of evidence indicates that non-antibiotic stressors can contribute to AMR evolution. Cross-resistance and co-selection mechanisms suggest that tolerance to one compound may predispose bacteria to acquire resistance to others, ultimately promoting multidrug resistance strains [15]. Environmental contaminants such as plastics and toxic metals have been shown to enhance the dissemination of AMR genes [16–18], while biocides can similarly select for resistance through overlapping of linked mechanisms [19]. In contrast, the role of pesticides in antimicrobial resistance has only recently begun to receive attention [20–22].

Glyphosate, the most widely used herbicide globally, represents a particularly relevant yet underexplored selective pressure. By targeting 5-enolpyruvylshikimate-3-phosphate synthase (EPSPS), glyphosate disrupts the shikimate pathway, a central metabolic route in many bacteria. Previous studies have demonstrated that glyphosate residues can alter microbial community composition and function in soil ecosystems, with downstream effects on nutrient-cycling and plant–soil processes [21, 22]. Our previous work has further characterized the molecular mechanisms underlying bacterial adaptation to glyphosate [23]. Importantly, recent evidence suggests that glyphosate exposure may also promote AMR, particularly in multidrug-resistant bacterial strains [24]. Although glyphosate is not mutagenic and acts *via* a mechanism distinct from that of antibiotics, co-exposure can induce efflux pump activity and, indirectly, facilitate the emergence of antibiotic resistance [25].

Despite these insights, it remains unclear whether intrinsic bacterial sensitivity to glyphosate systematically influences AMR gene content and its evolutionary dynamics across genomes. Addressing this gap is critical for understanding how widespread environmental chemicals contribute to the structuring of AMR genes in microbial communities. We therefore hypothesized that glyphosate sensitivity, determined by variation in the target enzyme EPSPS, shapes both the composition and evolutionary trajectories of AMR genes across diverse bacterial lineages.

## Material and methods summary

To test this hypothesis, we compiled a comprehensive dataset of bacterial genomes from the human microbiota available in the NCBI genome database [26] and environmental taxa from the Alignable Tight Genomic Clusters (ATGC) resource [27]. The glyphosate target enzyme, EPSPS, was identified using BLAST searches against the COG0128 orthologous group [28], and sequences were classified as glyphosate-sensitive, resistant, or unclassified using EPSPSClass webserver [28]. AMR genes and their associated resistance mechanisms were annotated with the Comprehensive Antibiotic Resistance Database (CARD) [29]. To characterize the evolutionary dynamics of AMR genes, we applied two complementary phylogenetic approaches. First, we used Count [30] to estimate rates of gene gain, loss, expansion, and contraction under a maximum likelihood birth-and-death model. Second, we employed Gloome [31] to infer gene-gain and gene-loss events along phylogenies using stochastic mapping. Finally, we evaluated the extent to which genomic features correlate with AMR gene content and modeled AMR accumulation across bacterial groups differing in EPSPS-based glyphosate sensitivity (see supplementary methods).

## Results

### Antimicrobial Resistance (AMR) diversity across EPSPS sensitivity classes

We compared AMR profiles across human-microbiota genomes and environmental taxa from the ATGC dataset by grouping species according to EPSPS-based glyphosate sensitivity (Tables S1–S4). Across all genomes, we identified resistance to 40 antibiotics (Tables S3–S4), mediated by six major AMR mechanisms, including antibiotic efflux, antibiotic inactivation, target alteration, target protection, target replacement, and reduced membrane permeability (Tables S1–S2). Individual species frequently encoded multiple resistance mechanisms simultaneously. A consistent pattern emerged across both datasets: EPSPS-sensitive bacteria harbored a greater diversity of AMR determinants than EPSPS-resistant bacteria (Table 1). In the human microbiota, this difference was moderate, with sensitive-to-resistant (S/R) ratios ranging from 1.35 to 1.72 (drugs) and 1.32 to 1.68 (mechanisms). In contrast, the ATGC dataset showed substantially larger differences, with S/R ratios ranging between 14.14 to 29.94 (drugs) and 9.23 and 19.55. These results indicate that glyphosate-sensitive bacteria, particularly environmental taxa, harbor a broader repertoire of AMR genes and mechanisms than glyphosate-resistant strains.

**Table 1.** Number of antimicrobial resistant (AMR) drugs and mechanisms in glyphosate-sensitive and resistant bacteria.

| EPSPS | Drugs / Mbp | Mechanism / Mbp | Drugs / sp | Mechanisms / sp |
| --- | --- | --- | --- | --- |
| **Human Microbiota dataset** |  |  |  |  |
| Sensitive (S) | 7.27 | 2.64 | 30.55 | 11.08 |
| Resistant (R) | 5.37 | 1.99 | 17.7 | 6.6 |
| S/R | **1.35** | **1.32** | **1.72** | **1.68** |
| **Alignable Tight Genomic Clusters (ATGC) dataset** |  |  |  |  |
| Sensitive (S) | 3.49 | 1.33 | 13.87 | 5.31 |
| Resistant (R) | 0.25 | 0.14 | 0.46 | 0.27 |
| S/R | **14.14** | **9.23** | **29.94** | **19.55** |

*ATGC: Alignable Tight Genomic Clusters, AMR: Antimicrobial Resistance, Mbp: Megabase pairs, Drugs/Mbp: Number of AMR drugs per Mbp, Mechanisms/Mbp: Number of AMR mechanism / Mbp, Drugs/sp: Number of AMR drugs per species, Mechanisms/sp: Number of AMR mechanisms per species*

### Antimicrobial Resistance (AMR) accumulation patterns are decoupled from genome features

To determine whether these differences reflect underlying genome architecture, we tested for associations between genomic features (genome length, gene number, protein number, and GC content) and AMR abundance within each EPSPS class. Neither Pearson nor Spearman analyses revealed significant correlations in either dataset (Figures S1–S3), indicating that AMR content is largely independent of genome structure. In contrast, probabilistic analyses revealed clear differences between EPSPS classes. Cumulative probability distribution and Poisson models showed that EPSPS-resistant bacteria accumulated AMR genes (including both drug resistance genes and resistance mechanisms) at higher rates than EPSPS-sensitive bacteria in both datasets (Figure S4). Although the human microbiota genomes contained more AMR elements overall, the relative accumulation rates were consistent across datasets. Together, these results demonstrate that while total AMR burden varies across taxa, the rate of AMR acquisition is strongly associated with EPSPS sensitivity rather than genome architecture.

### Elevated evolutionary turnover of AMR genes in glyphosate-resistant bacteria

To further resolve the evolutionary processes underlying these patterns, we quantified AMR gene gain and loss dynamics using complementary phylogenetic approaches. Analyses using Count and Gloome consistently showed that EPSPS-resistant bacteria exhibit higher rates of AMR gene gain, loss, expansion, and reduction than sensitive bacteria (Figure 1; Tables S5–S6). These results contrast with the observed distribution of AMR content: although EPSPS-sensitive bacteria harbor more AMR genes, EPSPS-resistant taxa exhibit markedly higher rates of gene turnover. Probabilistic Poisson-based analyses corroborated this, indicating faster accumulation in resistant lineages for both AMR drug classes and resistance mechanisms.

**Figure 1.**
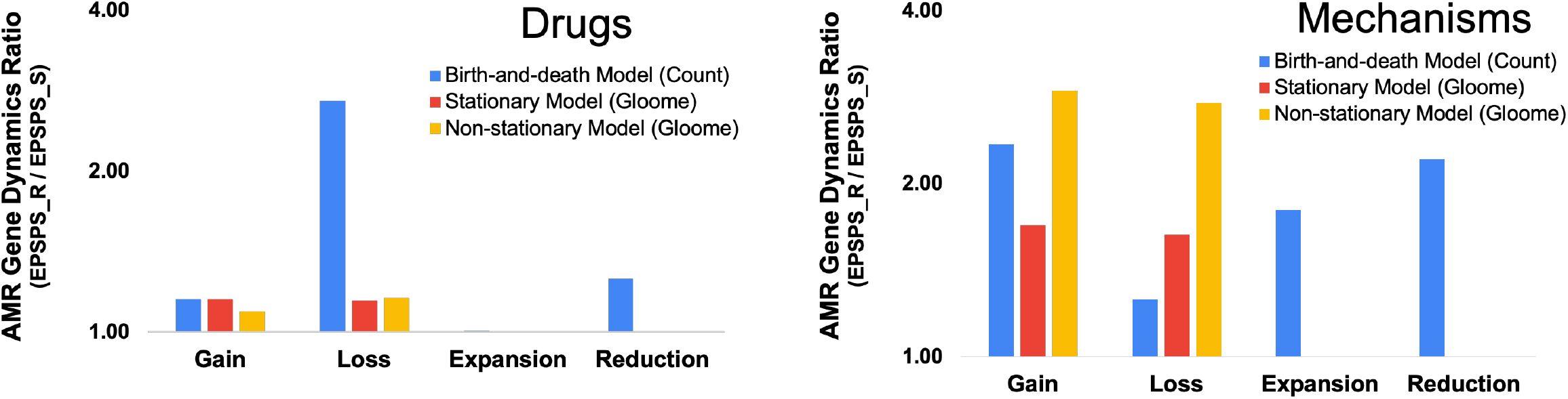
Ratio of the antimicrobial resistance (AMR; drugs and mechanisms) dynamics (gain, loss, expansions, and reductions) in EPSPS resistant (EPSPS_R) and sensitive (EPSPS_S) bacteria in the alignable tight genomic clusters (ATGC). The figure shows the result of the three independent models Count’s birth-and-death model [30] and Gloome’s stationary and none-stationary models [31].

Together, these findings reveal a key distinction between AMR content and AMR dynamics. Glyphosate-sensitive bacteria tend to maintain a larger reservoir of AMR genes, whereas glyphosate-resistant bacteria exhibit accelerated evolutionary flux of these genes. This suggests that glyphosate resistance is associated not with higher AMR load per se, but with increased rates of AMR gene turnover, potentially enhancing the capacity of resistant lineages to rapidly adapt to antimicrobial pressures.

## Discussion

Glyphosate is known to shape microbial community composition and function [22], and our findings extend this view by showing that glyphosate sensitivity is closely associated with the evolutionary dynamics of antimicrobial resistance (AMR). While glyphosate-sensitive bacteria harbor a greater overall repertoire of AMR genes, glyphosate-resistant strains exhibit significantly higher rates of AMR gene turnover, including elevated rates of gene gain, loss, expansion, and reduction. This is consistently supported by both phylogenetic and probabilistic analyses.

These findings reveal a decoupling between AMR content and AMR dynamics. Sensitive taxa maintain a larger pool of AMR genes, whereas resistant taxa display more evolutionarily dynamic genomes. Both modeling approaches indicate that the acquisition of glyphosate resistance is associated with increased genomic fluidity, which may enhance bacteria’s capacity to respond rapidly to fluctuating environmental pressures and antimicrobial challenges. A plausible mechanistic basis for this lies in the overlap between tolerance pathways to glyphosate and antibiotics. In particular, efflux-based mechanisms are known to mediate resistance to a wide range of compounds, and co-exposure experiments have shown that glyphosate can modulate bacterial susceptibility to specific antibiotics [23,33]. Such shared or linked resistance pathways provide a route for co-selection, whereby adaptation to one stressor indirectly promotes resistance to others.

Importantly, this interaction may be especially relevant in the context of human and livestock-associated microbiomes. Antibiotics are widely used in clinical and veterinary settings, while glyphosate is extensively applied in crop and fodder production, creating multiple opportunities for indirect human and livestock exposure through diet and the environment. Residues entering the food chain or drinking water can influence human and livestock-associated microbial communities, including the gut microbiota, where they may act as selective agents. More broadly, given that many herbicides and other agrochemicals influence host-associated microbiomes [22], our findings likely extend beyond glyphosate. This underscores the need to incorporate pesticide-driven selection into AMR risk frameworks and to adopt a more integrated perspective that links environmental exposures to human health outcomes.

## Conclusions

Our results indicate that glyphosate resistance is linked to increased AMR gene turnover rather than higher AMR gene abundance. Elevated rates of AMR gene gain, loss, expansion and reduction suggest enhanced genomic plasticity in glyphosate-resistant bacteria, increasing the selective availability of resistance mechanisms. Furthermore, this accelerated turnover may facilitate the fixation of advantageous AMR traits (i.e. increasing the AMR mechanisms and the resistance against a wider range for drugs) under favorable selective conditions, thereby promoting the emergence of multidrug-resistant lineages.

## Authors contributions

Conceptualization: PP. Critical revision: TT, SAM, MH, KS, IS, JI, PP. Methodology: TT, JI, EV, PP. Data collection: TT, LL, MJR, PP. Computational model: TT, EV, JI, PP. Formal investigation: TT, JI, PP. Resources, MH, KS, JI, PP, TT. Supervision: PP. Writing—original draft preparation: TT, PP. Writing—review and editing TT, JI, SAM, MH, KS. Writing the final version of the manuscript: PP. All authors have read and agreed to the published version of the manuscript.

## Acknowledgments

This research was funded by the Academy of Finland (grant no. 311077 to Marjo Helander), and the Jenny and Antti Wihuri Foundation (to Tuomas Tall).

