## Supplementary material for "Glyphosate as a driver of antimicrobial resistance evolution in bacteria"

### SUPPLEMENTARY METHODS

*Human microbiome*

We have analyzed 5800 bacterial genomes from the human microbiome. The genomes were gathered from the NCBI Datasets: Genomes [1]. Genomes were mapped through BLAST searches onto the COG0128 from the database of Cluster of Orthologous Groups (COG) to identify EPSPS proteins [2].

*Dataset of alignable tight genomic clusters (ATGC)*

Proteins were gathered from the Alignable Tight Genomic Clusters (ATGC) database, which encompasses over 4.5 million protein sequences from over 1,500 prokaryotic genomes that pass the criteria of exemplar ATGCs [3], as described in Puigbò et al. 2014 [4]. BLAST annotated EPSPS protein sequences were added onto the Clusters of Orthologous Genes (COG) database [2]. We analysed 3074 bacterial ATGCs and selected clusters with more than 5 species to further analyses.

*Detection of resistance to glyphosate*

The herbicide glyphosate works by blocking the cell's shikimate pathway by inhibiting the 5-enolpyruvylshikimate-3-phosphate synthase enzyme (EPSPS) [5]. EPSPS enzyme's potential sensitivity to glyphosate is assessed in the EPSPSClass server (https://ppuigbo.me/programs/EPSPSClass/) [6]. EPSPS can be classified based on known amino acid markers and motifs [7–12]. The EPSPS classes are class I (alpha or beta) [11, 12], class II [7, 8], class III [9] and class IV [10]. However, a great number of microbes are still unclassified and so their potential glyphosate sensitivity is unknown [6]. The four reference sequences that are used to distinguish the EPSPS classes are from *Vibrio cholerae serotype O1* (vcEPSPS, class I), *Coxiella burnetii* (cbEPSPS, class II), *Brevundimonas vesicularis* (bvEPSPS, class III) and *Streptomyces davawensis* (sdEPSPS, class IV).

*Antibiotic resistance genes with CARD*

The identification of antibiotic resistant genes was determined based on the Comprehensive Antibiotic Resistance Database (CARD) analysis [13]. The CARD is a bioinformatic library of antibiotic resistance with curated antimicrobial resistance genes (ARG) sequences. We have analyzed 5799 bacterial strains from the human microbiome dataset and 3703 bacterial groups from the ATGC database. We have analyzed 10 individuals and a combination of mechanisms of antibiotic resistance including antibiotic efflux, antibiotic inactivation, antibiotic target alteration and reduced permeability to antibiotics (Individual AR mechanisms number at 6) (Table S1-4).

*Birth and death model of antimicrobial resistance*

The analysis of the family history of posterior probabilities was done using the Count program. The Count program is a software package for the analysis of numerical profiles on a phylogeny. The Program was used to estimate the gain, loss, expansion, and reduction rates of antibiotic genes and the combination of genes [4, 14, 15]. To calculate the rates, Count needs two inputs: a matrix that contains the number of gene copies, and a rooted species tree. The gene copy matrix was obtained from the CARD (i.e. two matrices were created: 1) total number of genes resistant to specific drug, and 2) total number of gene copies for an antimicrobial resistance mechanism), while the rooted family tree was obtained from the ATGC database. The program calculates these rates using a phylogenetic birth-and-death model that requires the following parameters: $\kappa$ (rate of gene gain), $\lambda$

(individual gene duplication rate) and μ (individual gene loss rate) X. Thus, the parameters (κ, λ, μ) are different for each gene family and across edges of the species tree. These parameters are computed by Count using ML optimization. It is recommended that the parameters are optimized iteratively, in several rounds of increasing computational complexity, such that in each round the rates from the previous round are used as the starting point. We optimized the four parameters through four rounds of increasing complexity. The parameter values obtained in the final round were used to estimate the numbers of gains, losses, expansions and reductions for all gene families at different branches of the species tree (Table S6). This final analysis was performed using the `posteriors' option of Count, which analyses and integrates several phylogenetic scenarios and calculates rates of gain, loss, expansion and reduction across all branches. The sum across all branches and across all families is taken as the estimate of the number of events across the entire history of a given group of organisms. The sum was calculated with the use of custom code that ran through all analysed samples. There were matrices and trees that could not be analysed, leaving 189 posterior analyses to be summed [4, 14, 15].

*Inference of gene gain and loss via stochastic mapping*

Gene gains and losses at each branch of each species tree were estimated with the standalone version of the gain-loss mapping engine Gloome [16]. As inputs, we used the same rooted species trees as those provided to Count and a set of gene presence/absence matrices (phyletic profiles) derived from the copy number matrices described above and reformatted as required by Gloome. The parameter configuration file was set to optimize the likelihood of the observed phyletic profiles under a genome evolution model with 4 categories of gamma-distributed gain and loss rates. For each dataset, we ran Gloome in two modes: (i) assuming stationary state, with frequencies at the root equal to those in equilibrium; and (ii) allowing for out-of-equilibrium (non-stationary) dynamics, with equal frequencies at the root for all genes. Unlike Count, Gloome does not account for gene copy number and, consequently, it does not estimate expansion and reduction rates.

*Statistical analysis*

Spearman and Pearson analyses were done with a Python program with spicy.stats.spearmanr and.pearsonr modules. Analysis was done on ATGC species. To remove noise from analysis we calculated the average values of DNA lengths, total number of genes, total number of proteins and total GC content of DNA, and average number of AMR drugs and AMR mechanisms. To further limit noise we selected ATGCs that had five or more subspecies [17–20].

Using these averages we calculated the Spearman and Pearson correlation and p-values separately for EPSPS resistant, sensitive and unclassified species. We compared the average values of DNA lengths, total number of genes, total number of proteins and total GC content of DNA to the average number of AMR drugs and AMR mechanisms. The significant p-value was 0.05 (Figure S2).

We calculated the values for species that did have AMR drugs or mechanisms and for all species even if they didn't have those. The values of ATGC:s that had AMR genes were published in this paper. In addition, we used Python to make scatter plots of each value pair (Figure S1).

*Poisson distribution*

We used a Poisson distribution to model the probability that bacteria accumulate a given number of AMR elements. The Poisson model is appropriate for discrete count data and describes the probability of observing k events within a defined interval of evolutionary time or lineage space. It is defined by a single parameter, λ (lambda), which represents the mean rate at which events occur. In this context, λ corresponds to the expected number of AMR gene acquisitions per bacterial lineage. Using λ, we estimated the cumulative probability of AMR gain across EPSPS sensitivity classes and compared whether glyphosate-resistant and glyphosate-sensitive bacteria differ in their expected rates of AMR accumulation.

### SUPPLEMENTARY FIGURES

**a)** **b)**

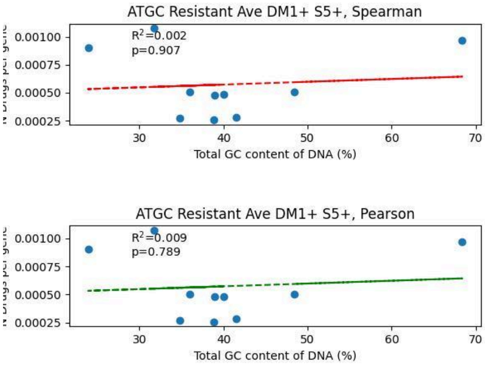


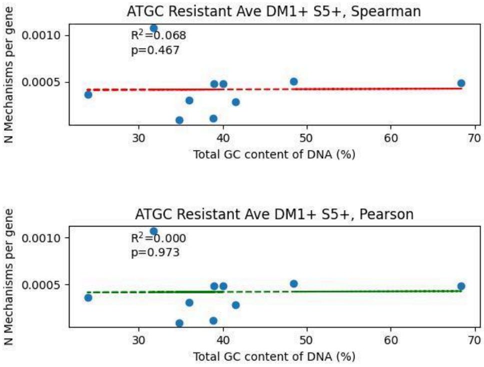


**c)** **d)**

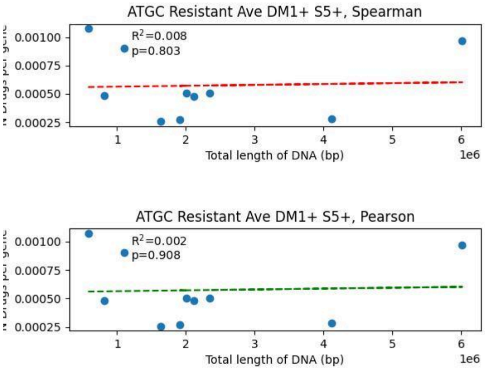


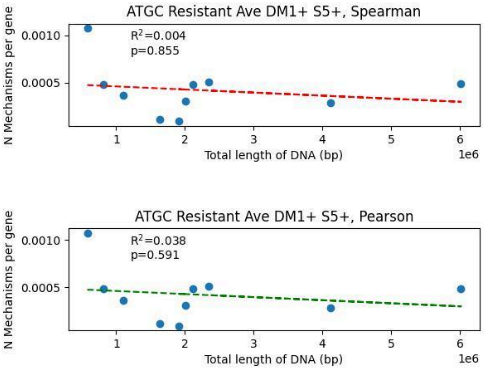


**e)** **f)**

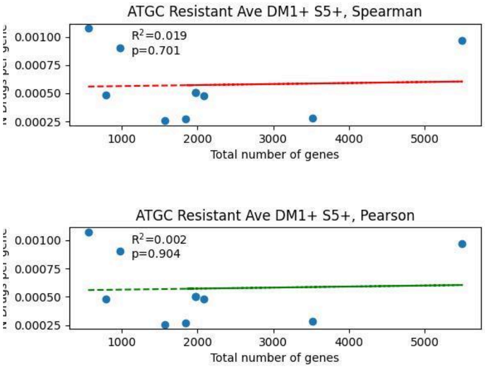


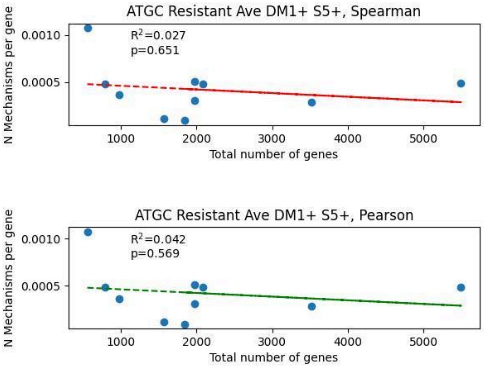


**g)** **h)**

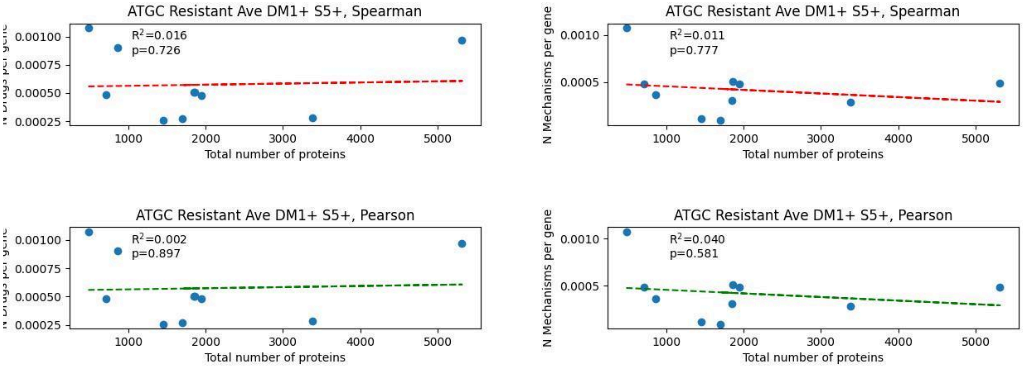


**Figure S1.** Scatter plots of correlation between ATGC's genomics parameters and Antibiotic resistance parameters in EPSPS-resistant species. ATGC species that had resistant EPSPS gene and more than 5 species per ATGC and had antibiotic resistance. a) Total CG presence in DNA (%) / Total number of antibiotic drugs resisted per gene, b) Total CG presence in DNA (%) / Total number of antibiotic resistance mechanisms per gene, c) Total length of DNA (bp) / Total number of antibiotic drugs resisted per gene, d) Total length of DNA (bp) / Total number of antibiotic resistance mechanisms per gene, e) Total number of Genes / Total number of antibiotic drugs resisted per gene, f) Total number of Genes / Total number of antibiotic resistance mechanisms per gene, g) Total number of proteins / Total number of antibiotic drugs resisted per gene and h) Total number of proteins / Total number of antibiotic resistance mechanisms per gene

**a)** **b)**

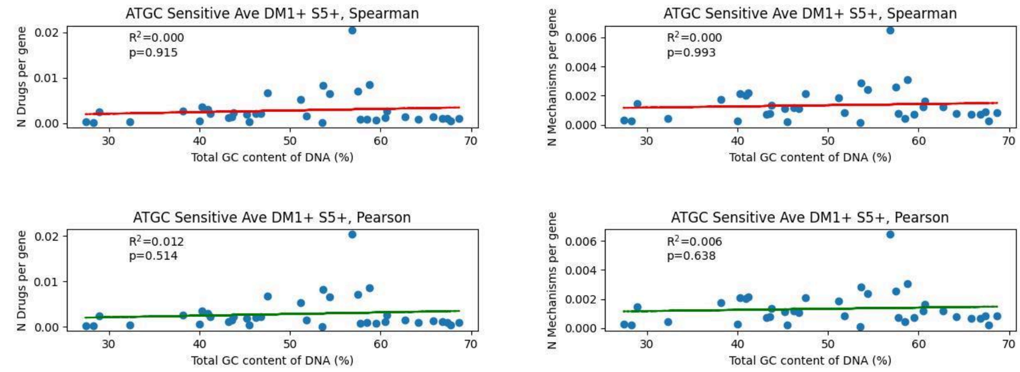


**c)** **d)**

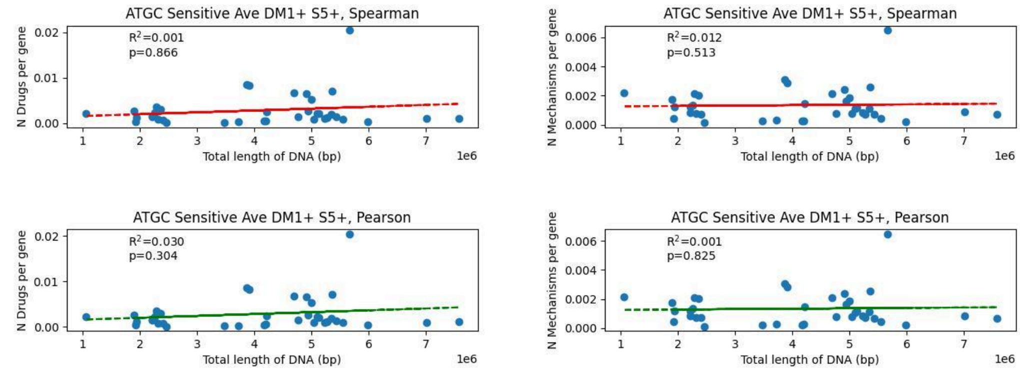


**e)** **f)**

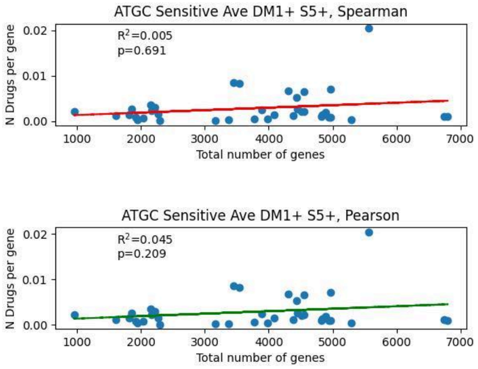


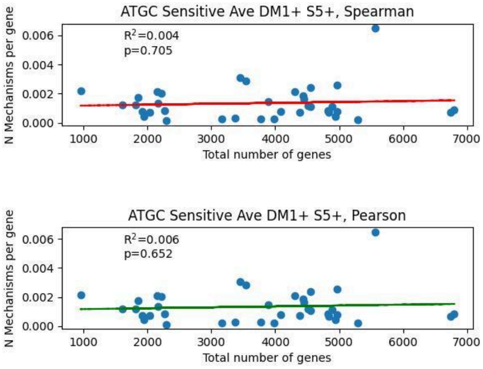


**g)** **h)**

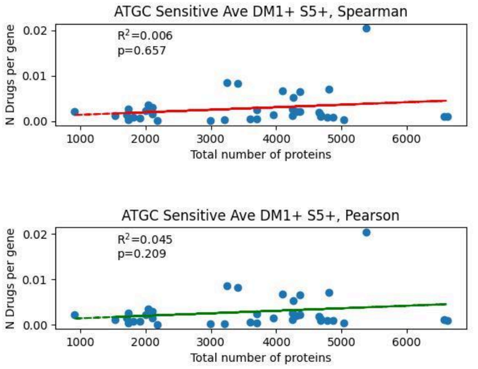


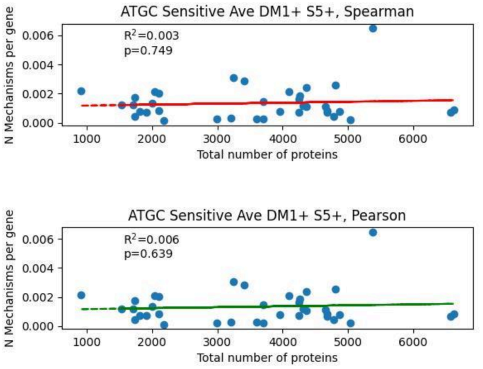


**Figure S2.** Scatter plots of correlation between ATGC's genomics parameters and Antibiotic resistance parameters in EPSPS Sensitive species. ATGC species that have sensitive EPSPS gene and more than 5 species per ATGC and had antibiotic resistance. a) Total CG presence in DNA (%) / Total number of antibiotic drugs resisted per gene, b) Total CG presence in DNA (%) / Total number of antibiotic resistance mechanisms per gene, c) Total length of DNA (bp) / Total number of antibiotic drugs resisted per gene, d) Total length of DNA (bp) / Total number of antibiotic resistance mechanisms per gene, e) Total number of Genes / Total number of antibiotic drugs resisted per gene, f) Total number of Genes / Total number of antibiotic resistance mechanisms per gene, g) Total number of proteins / Total number of antibiotic drugs resisted per gene and h) Total number of proteins / Total number of antibiotic resistance mechanisms per gene.

**a)** **b)**

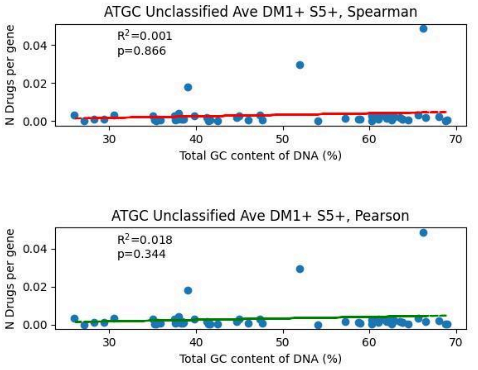


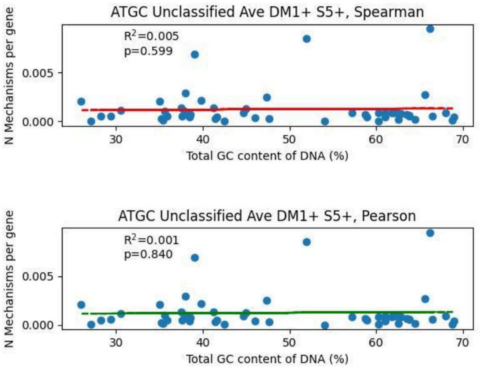


**c)** **d)**

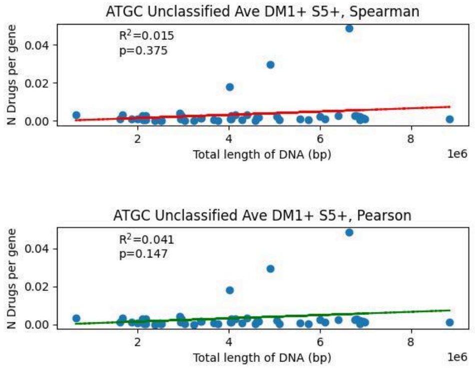


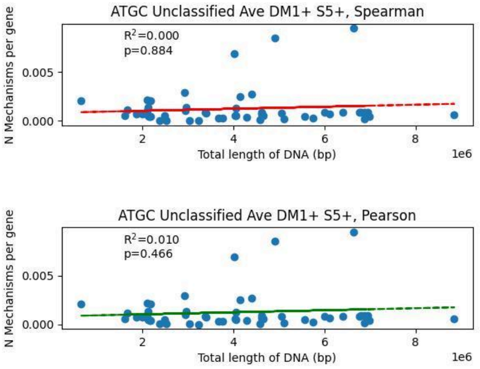


**e)**

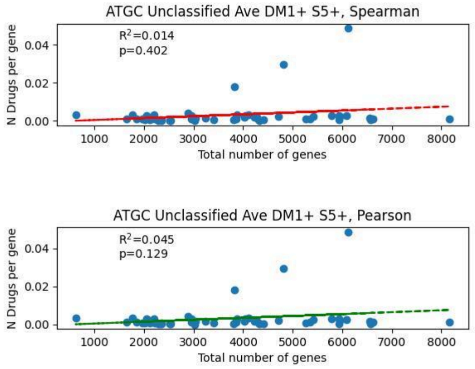


**f)**

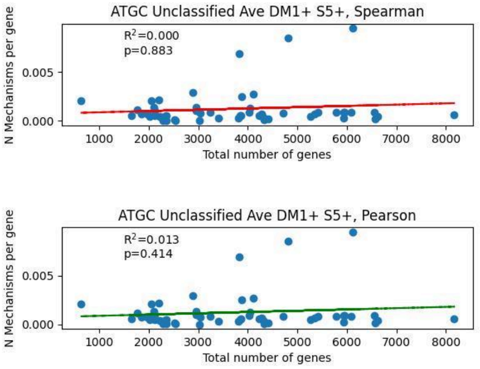


**g)**

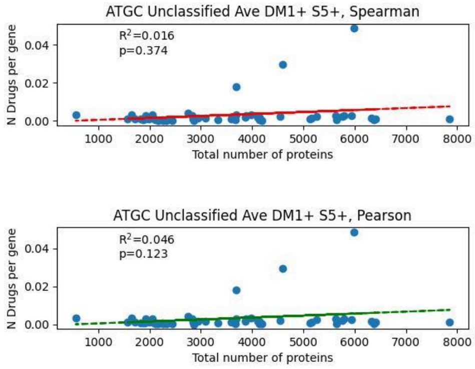


**h)**

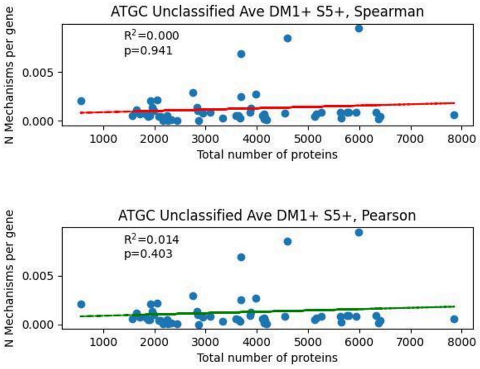


**Figure S3.** Scatter plots of correlation between ATGC's genomics parameters and Antibiotic resistance parameters in EPSPS Unclassified species. ATGC species that have unclassified EPSPS gene and more than 5 species per ATGC and had antibiotic resistance. a) Total CG presence in DNA (%) / Total number of antibiotic drugs resisted per gene, b) Total CG presence in DNA (%) / Total number of antibiotic resistance mechanisms per gene, c) Total length of DNA (bp) / Total number of antibiotic drugs resisted per gene, d) Total length of DNA (bp) / Total number of antibiotic resistance mechanisms per gene, e) Total number of Genes / Total number of antibiotic drugs resisted per gene, f) Total number of Genes / Total number of antibiotic resistance mechanisms per gene, g) Total number of proteins / Total number of antibiotic drugs resisted per gene and h) Total number of proteins / Total number of antibiotic resistance mechanisms per gene.

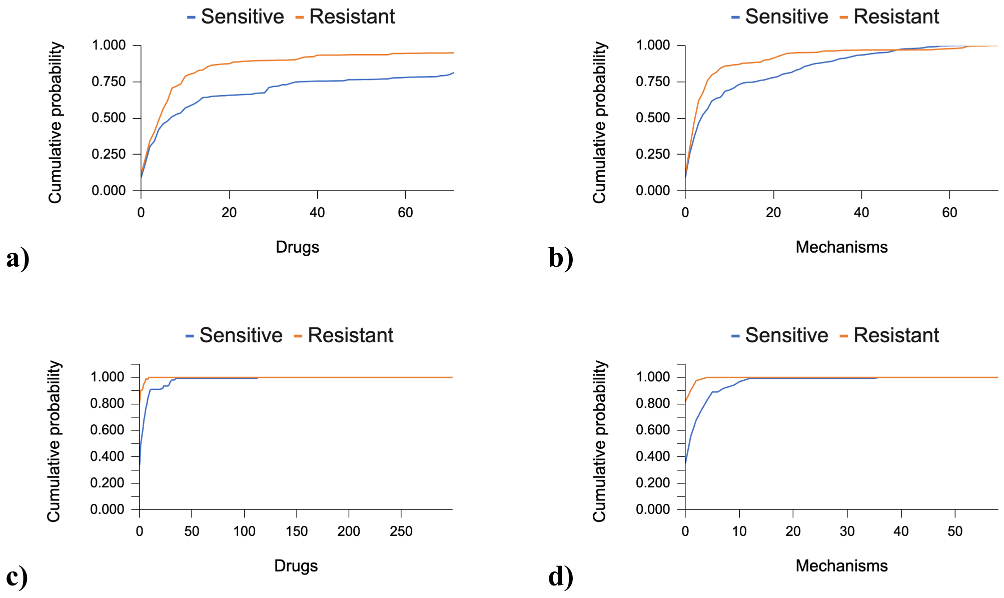


**Figure S4.** Probability of gaining AMR drugs and mechanisms in the human microbiota and the alignable tight genomic clusters
a) Cumulative probability that EPSPS sensitive or resistant ATGC species gain AMR drug genes.
b) Cumulative probability that EPSPS-sensitive or resistant ATGC species gain AMR mechanisms.
c) Cumulative probability that EPSPS-sensitive or resistant human microbiota species gain AMR drug genes.
d) Cumulative probability that EPSPS-sensitive or resistant human microbiota species gain AMR drug mechanisms.

### SUPPLEMENTARY TABLES

**Table S1.** Antibiotic Resistance Mechanisms in bacteria from ATGC. Total number of bacteria and number of bacteria that's EPSP synthase gene is sensitive to glyphosate, gene's sensitivity is unclassified or gene is absent) and bacteria whose genes are resistant to herbicides.

| Mechanism | Total | EPSPS Sensitive | EPSPS Unclassified | EPSPS Resistant |
|---|---|---|---|---|
| Antibiotic efflux | 18892 | 13802 | 192 | 4898 |
| Antibiotic efflux; Reduced permeability to antibiotics | 614 | 498 | 5 | 111 |
| Antibiotic inactivation | 4402 | 2755 | 46 | 1601 |
| Antibiotic target alteration | 5984 | 5008 | 89 | 887 |
| Antibiotic target alteration; Antibiotic efflux | 1486 | 1104 | 12 | 370 |
| Antibiotic target alteration; Antibiotic efflux; Reduced permeability to antibiotic | 388 | 385 | 3 | 0 |
| Antibiotic target alteration; Antibiotic target replacement | 130 | 119 | 2 | 9 |
| Antibiotic target protection | 471 | 265 | 18 | 188 |
| Antibiotic target replacement | 593 | 352 | 6 | 235 |
| Reduced permeability to antibiotic | 284 | 210 | 2 | 72 |

**Table S2.** Antibiotic Resistance Mechanisms in bacteria from Human Microbiota. Total number of bacteria and number of bacteria that's *epsp synthase* gene is sensitive to glyphosate, gene's sensitivity is unclassified or gene is absent) and bacteria that's gene is resistant to herbicide.

| Mechanism | Total | EPSPS Sensitive | EPSPS Unclassified | EPSPS Resistance |
|---|---|---|---|---|
| Antibiotic efflux | 24419 | 16993 | 789 | 6637 |
| Antibiotic efflux; Reduced permeability to antibiotic | 866 | 726 | 8 | 132 |
| Antibiotic inactivation | 10219 | 6123 | 265 | 3831 |
| Antibiotic target alteration | 9791 | 7720 | 116 | 1955 |
| Antibiotic target alteration; Antibiotic efflux | 1656 | 1187 | 15 | 454 |
| Antibiotic target alteration; Antibiotic efflux; Reduced permeability to antibiotic | 334 | 334 | 0 | 0 |
| Antibiotic target alteration; Antibiotic target replacement | 213 | 174 | 0 | 39 |
| Antibiotic target protection | 1340 | 790 | 17 | 533 |
| Antibiotic target replacement | 1839 | 1142 | 22 | 675 |
| Reduced permeability to antibiotic | 651 | 427 | 126 | 98 |

**Table S3.** Number of antibiotic resistant bacteria in ATGC by their specific antibiotic. Total number of bacteria and number of bacteria that's *epsp synthase* gene is sensitive to glyphosate, gene's sensitivity is unclassified or gene is absent) and bacteria that's gene is resistant to herbicide.

| Drug | Total | EPSPS Sensitive | EPSPS Unclassified | EPSPS Resistant |
|---|---|---|---|---|
| Acridinedye | 1891 | 670 | 18 | 1203 |
| Aminocoumarinantibiotic | 2413 | 1518 | 24 | 871 |
| Aminoglycosideantibiotic | 5277 | 3935 | 52 | 1290 |
| Antibacterialfreefattyacids | 27 | 27 | 0 | 0 |
| Benzalkoniumchloride | 404 | 345 | 3 | 56 |
| Bicyclomycin | 55 | 0 | 1 | 54 |
| Carbapenem | 3046 | 2008 | 30 | 1008 |
| Cephalosporin | 9859 | 8039 | 78 | 1742 |
| Cephamycin | 3874 | 3219 | 35 | 620 |
| Diaminopyrimidineantibiotic | 2324 | 905 | 30 | 1389 |
| Elfamycinantibiotic | 1477 | 1457 | 16 | 4 |
| Ethionamide | 2 | 2 | 0 | 0 |
| Fluoroquinoloneantibiotic | 13728 | 9900 | 148 | 3680 |
| Fosfomycin | 1533 | 1150 | 8 | 375 |
| Fusidicacid | 108 | 1 | 1 | 106 |
| Glycopeptideantibiotic | 116 | 29 | 1 | 86 |
| Glycylcycline | 3730 | 3322 | 29 | 379 |
| Isoniazid | 120 | 116 | 4 | 0 |
| Lincosamideantibiotic | 748 | 223 | 8 | 517 |
| Macrolideantibiotic | 6495 | 4179 | 79 | 2237 |
| Monobactam | 2355 | 1531 | 29 | 795 |

| | | | | |
|---|---|---|---|---|
| Mupirocin | 11 | 5 | 0 | 6 |
| Nitrofuranantibiotic | 128 | 126 | 2 | 0 |
| Nitroimidazoleantibiotic | 507 | 495 | 12 | 0 |
| Nucleosideantibiotic | 766 | 674 | 5 | 87 |
| Oxazolidinoneantibiotic | 230 | 20 | 1 | 209 |
| Para-Aminosalicylicacid | 17 | 17 | 0 | 0 |
| Penam | 10336 | 8607 | 83 | 1646 |
| Penem | 2629 | 1934 | 26 | 669 |
| Peptideantibiotic | 4019 | 2980 | 36 | 1003 |
| Phenicolantibiotic | 6710 | 4410 | 66 | 2234 |
| Pleuromutilinantibiotic | 137 | 18 | 12 | 107 |
| Polyamineantibiotic | 50 | 50 | 0 | 0 |
| Pyrazinamide | 15 | 15 | 0 | 0 |
| Rhodamine | 347 | 345 | 2 | 0 |
| Rifamycinantibiotic | 4948 | 4587 | 37 | 324 |
| Streptograminantibiotic | 310 | 56 | 7 | 247 |
| Sulfonamideantibiotic | 727 | 247 | 9 | 471 |
| Tetracyclineantibiotic | 11155 | 7516 | 151 | 3488 |
| Triclosan | 3793 | 3206 | 32 | 555 |

**Table S4.** Number of Antibiotic Resistant Bacteria in human microbiota. Total number of bacteria and number of bacteria that's *epsp synthase* gene is sensitive to glyphosate, gene's sensitivity is unclassified or gene is absent) and bacteria that's gene is resistant to herbicide.

| Drug | Total | EPSPS Sensitive | EPSPS Unclassified | EPSPS Resistance |
|---|---|---|---|---|
| Acridinedye | 1754 | 281 | 3 | 1470 |
| Aminocoumarinantibiotic | 2625 | 1439 | 8 | 1178 |
| Aminoglycosideantibiotic | 8531 | 6046 | 291 | 2194 |
| Antibacterialfreefattyacids | 57 | 57 | 0 | 0 |
| Benzalkoniumchloride | 347 | 281 | 0 | 66 |
| Bicyclomycin | 66 | 0 | 0 | 66 |
| Carbapenem | 4836 | 3161 | 264 | 1411 |
| Cephalosporin | 13816 | 10637 | 349 | 2830 |
| Cephamycin | 5301 | 4276 | 152 | 873 |
| Diaminopyrimidineantibiotic | 3762 | 1863 | 26 | 1873 |
| Elfamycinantibiotic | 2677 | 2659 | 18 | 0 |
| Ethionamide | 3 | 3 | 0 | 0 |
| Fluoroquinoloneantibiotic | 19175 | 13456 | 557 | 5162 |
| Fosfomycin | 2320 | 1673 | 19 | 628 |
| Fusidicacid | 204 | 0 | 1 | 203 |
| Glycopeptideantibiotic | 619 | 159 | 2 | 458 |
| Glycylcycline | 3764 | 3294 | 32 | 438 |
| Isoniazid | 66 | 66 | 0 | 0 |
| Lincosamideantibiotic | 1238 | 336 | 9 | 893 |
| Macrolideantibiotic | 9083 | 5696 | 66 | 3321 |

| | | | | |
|---|---|---|---|---|
| Monobactam | 3192 | 2075 | 139 | 978 |
| Mupirocin | 13 | 5 | 0 | 8 |
| Nitrofuranantibiotic | 411 | 403 | 8 | 0 |
| Nitroimidazoleantibiotic | 759 | 716 | 43 | 0 |
| Nucleosideantibiotic | 623 | 337 | 0 | 286 |
| Oxazolidinoneantibiotic | 344 | 93 | 4 | 247 |
| Para-Aminosalicylicacid | 19 | 19 | 0 | 0 |
| Penam | 13491 | 10414 | 401 | 2676 |
| Penem | 3867 | 2777 | 153 | 937 |
| Peptideantibiotic | 5206 | 3810 | 97 | 1299 |
| Phenicolantibiotic | 7394 | 4679 | 40 | 2675 |
| Pleuromutilinantibiotic | 202 | 93 | 3 | 106 |
| Polyamineantibiotic | 45 | 45 | 0 | 0 |
| Pyrazinamide | 18 | 18 | 0 | 0 |
| Rhodamine | 281 | 281 | 0 | 0 |
| Rifamycinantibiotic | 5604 | 5056 | 41 | 507 |
| Streptograminantibiotic | 652 | 177 | 9 | 466 |
| Sulfonamideantibiotic | 1516 | 790 | 12 | 714 |
| Tetracyclineantibiotic | 15673 | 10489 | 531 | 4653 |
| Triclosan | 3551 | 2911 | 22 | 618 |

**Table S5. Models of gain, loss, expansion and reduction rates of gene copies of AMR genes and mechanisms of AMR in bacteria that have EPSPS gene(s) phylogenetic trees.**

| | | | | | | | | | | |
|---|---|---|---|---|---|---|---|---|---|---|
| **AMR Drugs (Birth-and-death model - Count)** | | | | | | | | | | |
| **EPSPS** | **Gain** | **Loss** | **Expansion** | **Reduction** | **N** | **Gain/N** | **Loss/N** | **Exp/N** | **Red/N** | **G/L** |
| Sensitive (S) | 221.2 | 56.8 | 89.5 | 31.8 | **39** | 5.67 | 1.46 | 2.30 | 0.81 | **3.90** |
| Resistant (R) | 306.9 | 184.9 | 108.5 | 48.2 | **47** | 6.53 | 3.93 | 2.31 | 1.03 | **1.66** |
| | | | | | **R/S** | **1.15** | **2.70** | **1.01** | **1.26** | |
| **AMR Mechanisms (Birth-and-death model - Count)** | | | | | | | | | | |
| **EPSPS** | **Gain** | **Loss** | **Expansion** | **Reduction** | **N** | **Gain/N** | **Loss/N** | **Exp/N** | **Red/N** | **G/L** |
| Sensitive (S) | 65.1 | 28.7 | 17.2 | 8.9 | **40** | 1.63 | 0.72 | 0.43 | 0.22 | **2.27** |
| Resistant (R) | 186.5 | 44.3 | 37.8 | 24.1 | **49** | 3.81 | 0.90 | 0.77 | 0.49 | **4.21** |
| | | | | | **R/S** | **2.34** | **1.26** | **1.80** | **2.21** | |
| **AMR Drugs (Stationary Model - Gloome)** | | | | | | | | | | |
| **EPSPS** | **Gain** | **Loss** | | | **N** | **Gain/N** | **Loss/N** | **Exp/N** | **Red/N** | **G/L** |
| Sensitive (S) | 2175 | 2295 | | | **42** | 51.8 | 54.7 | - | - | **0.95** |
| Resistant (R) | 2444 | 2565 | | | **41** | 59.6 | 62.6 | - | - | **0.95** |
| | | | | | **R/S** | **1.15** | **1.14** | **-** | **-** | |
| **AMR Mechanisms (Stationary Model - Gloome)** | | | | | | | | | | |
| **EPSPS** | **Gain** | **Loss** | | | **N** | **Gain/N** | **Loss/N** | **Exp/N** | **Red/N** | **G/L** |
| Sensitive (S) | 342.2 | 359.0 | | | 42.0 | 8.1 | 8.5 | - | - | 1.0 |
| Resistant (R) | 564.6 | 571.3 | | | 41 | 13.8 | 13.9 | - | - | 0.99 |
| | | | | | **R/S** | **1.69** | **1.63** | **-** | **-** | |
| **AMR Drugs (Non-stationary Model - Gloome)** | | | | | | | | | | |
| **EPSPS** | **Gain** | **Loss** | | | **N** | **Gain/N** | **Loss/N** | **Exp/N** | **Red/N** | **G/L** |
| Sensitive (S) | 1890.6 | 1876.6 | | | **28** | 67.5 | 67.0 | - | - | 1.01 |
| Resistant (R) | 2727.7 | 2876.6 | | | **37** | 73.7 | 77.7 | - | - | 0.95 |
| | | | | | **R/S** | **1.09** | **1.16** | **-** | **-** | |
| **AMR Mechanisms (Non-stationary Model - Gloome)** | | | | | | | | | | |
| **EPSP** | **Gain** | **Loss** | | | **N** | **Gain/N** | **Loss/N** | **Exp/N** | **Red/N** | **G/L** |
| Sensitive (S) | 189.1 | 187.5 | | | 35 | 5.4 | **5.4** | - | - | 1.01 |
| Resistant (R) | 611.6 | 576.7 | | | 39 | 15.7 | **14.8** | - | - | 1.06 |
| | | | | | **R/S** | **2.90** | **2.76** | **-** | **-** | |

*Conditions: ATGC > 5 species ATGC >= 1 drug and >= 1 mechanism No error in count or Gloome*

ESPS shows if *epsp* is sensitive or resistant to glyphosate. Gain is the sum of gain rates. Loss is the sum of loss rates. Expansion is the sum of expansion rates. Reduction is the sum of reduction rates. N is the number of rooted family trees from ATGC. Gain/N is the gain rate per gene family. Loss/N is the loss rate per gene family. Exp/N is the expansion rate per gene family. Red/N is the reduction rates per gene family. G/L is the gain/loss ratio. R/S is resistance values compared to sensitivity values.

**Table S6.** Count parameters

| ROUNDS | -opt_rounds | -max_paralogues | -uniform_gain | -uniform_duplication | -gain_k | -loss_k | -duplication_k |
|---|---|---|---|---|---|---|---|
| Round 1 | 100 | - | TRUE | - | - | - | - |
| Round 2 | 100 | - | - | TRUE | - | - | - |
| Round 3 | 100 | 100 | - | - | 1 | - | - |
| Round 4 | 100 | 100 | – | - | - | 1 | - |
| Round 5 | 100 | 100 | - | - | - | - | 1 |
| Round 6 | 100 | 1000 | - | - | 2 | - | |
| Round 7 | 100 | 1000 | - | - | - | 2 | - |
| Round 8 | 100 | 1000 | - | - | - | - | 2 |
| Round 9 | 100 | 1000 | - | - | 4 | - | - |
| Round 10 | 100 | 1000 | - | - | - | 4 | - |
| Round 11 | 100 | 1000 | - | - | - | - | 4 |
